# Assessing the relationships between neurological and psychiatric diseases with astrocyte subtypes and psychotropic medications

**DOI:** 10.1101/2021.09.22.461367

**Authors:** Xiaolu Zhang, Alyssa Wolfinger, Rammohan Shukla, Anna Lundh, Xiaojun Wu, Mackenzie Abel, Robert E. McCullumsmith, Sinead M. O’Donovan

**Affiliations:** Department of Neurosciences, University of Toledo, Toledo, Ohio 43614; Promedica Neurosciences Institute, Toledo, Ohio

## Abstract

Astrocytes have many important functions in the brain, but their roles in CNS disorders and their responses to psychotropic medications are still being elucidated. In this study, we used gene enrichment analysis to assess the relationships between different astrocyte subtypes, neurological and psychiatric diseases, and psychotropic medications. We also carried out qPCR analyses and “look-up” studies to further assess the chronic effects of these drugs on astrocyte marker gene expression. Our bioinformatic analysis identified differential gene enrichment of different astrocyte subtypes in CNS disorders. The “common” astrocyte subtype was the most frequently enriched across disorders, but the highest level of enrichment was found in depression, supporting a role for astrocytes in this disorder. We also identified common enrichment of metabolic and signal transduction-related biological processes in astrocyte subtypes and CNS disorders. However, enrichment of different psychotropic medications, including antipsychotics, antidepressants, and mood stabilizers, was limited in astrocyte subtypes. These results were confirmed by “look-up” studies and qPCR analysis, which also reported little effect of common psychotropic medications on astrocyte marker gene expression, suggesting that astrocytes are not a primary target of these medications. Overall, this study provides a unique view of astrocyte subtypes and the effect of medications on astrocytes in disease, which will contribute to our understanding of their role in CNS disorders and offers insights into targeting astrocytes therapeutically.

## Introduction

Astrocytes play many important roles in the brain, from facilitating metabolism and cell signaling [1, 2], to responding to injury through regulating oxidative stress [3]. However, the role of astrocytes in the context of many neurological and psychiatric disorders is still poorly understood. While studies of postmortem brain broadly support perturbation of astrocytes in CNS disorders, the findings can prove challenging to interpret. For example, there is evidence of both increased [4–8] and decreased [7, 9–19] astrocyte marker expression, increased [6, 10] and decreased [20–26] astrocyte cell counts, changes in astrocyte morphology [7, 27–29], as well as no significant changes in astrocyte expression [27, 30–35] in disorders like schizophrenia and major depressive disorder (MDD). These findings also vary depending on the brain region studied [6, 7, 9, 11, 21, 28, 36–39] as well as the psychiatric disorder studied [16]. A potential cause for these conflicting findings may be due to the complex responses of astrocytes to injury; astrocytes can undergo remodeling that lead to a range of responses [40] from complete atrophy [41, 42] to a reactive state termed “reactive astrogliosis” [43, 44].

Reactive astrogliosis describes a spectrum of heterogenous changes in gene expression [45–47], cell morphology [46–51], and overall function including fluid and ion homeostasis [52, 53], oxidative stress response [54–56] and synapse formation [57, 58]. These changes can also range from reversible to permanent [59, 60]. A diverse set of mechanisms play a role in inducing astrogliosis [61], which can have a protective or pathological effect [62–64], depending on the subtype of astrocyte involved [43, 45], and the type of injury sustained [43, 48, 61]. In murine models of ischemic injury or LPS-induced infection, changes in astrocyte gene expression were identified in both a common set of genes and genes that were uniquely regulated by each type of injury [43]. There are also different “types” of astrocytes that can have potentially different responses to injury. These subtypes are classified based on cellular morphology (e.g., protoplasmic, fibrous, bushy), location (e.g., frontal cortex, hippocampus, cerebellum) and primary functions (e.g., metabolic, structural, signaling) [65]. However, our understanding of the different types of astrocytes in humans and their roles in CNS disorders is still limited [65, 66]. Although inclusive of this wide spectrum of phenotypic changes, the term “reactive astrogliosis” will be used here as an umbrella term to encompass the range of possible changes that occur in astrocytes as a result of neurological insult.

In addition to disease heterogeneity and methodological differences that contribute to the variable astrocyte findings in postmortem studies, psychotropic medications are hypothesized to be a significant confounding factor [65]. Antipsychotic use is associated with changes in brain volume in postmortem brain in schizophrenia subjects [67, 68] and in nonhuman primates administered antipsychotic drugs [69, 70]. Antidepressant use is associated with reduced astrocyte cell counts in rodent studies [71] and reduced astrocyte marker gene expression in postmortem studies [15] although other studies report no changes in these variables in postmortem studies [12, 23, 24, 26, 72]. Mood stabilizer use in bipolar disorder has also been associated with changes in astrocyte morphology and density in rodent studies [73]. Despite the extensive number of studies examining astrogliosis in postmortem and animal studies of neurological and psychiatric disorders, it is still unclear what effect these medications have on astrocyte expression and function in these disorders.

In this study we apply bioinformatic analysis of transcriptomic datasets to assess the relationship between different astrocyte subtypes and CNS disorders, and between astrocytes and psychotropic medications. Our goal is first, to explore how different subtypes of astrocytes are implicated in CNS disorders and then, to determine whether psychotropic medications are associated with molecular changes in astrocytes in disease.

## Materials and Methods

### Astrocyte subtypes gene-sets

The gene-sets for each transcriptomically distinct subtype of mouse astrocyte were downloaded from the supplementary data (https://holt-sc.glialab.org/sc/) of Batiuk et al’s. single-cell RNA sequencing astrocyte study [74]. These gene-sets include genes enriched in each of five subtypes of astrocytes (AST1-5) and genes expressed in at least 60% of astrocytes (common) identified in the study.

As described in Batiuk et al., AST1-3 appear to be mature astrocyte subtypes. AST1 is found in the subpial layer and hippocampus. AST2 has high expression in cortical layers 2/3 and 5, lower expression in cortical layers 1, 4, and 6, and negligible hippocampal expression. AST3 subtype is expressed throughout cortical layers and hippocampus. AST4 is found predominately in the subgranular zone of the hippocampus and may represent a progenitor or hippocampal neural stem cell population [74]. AST5 subtype expression overlaps substantially with AST4 and is hypothesized to represent an intermediate state between progenitor and mature astrocyte. AST5 shows enrichment in cortical layer 2/3 and 5, in the stratum lacunosum-moleculare and dentate gyrus of the hippocampus, and subpial layers.

### Disease-Disease similarity

Curated disease-associated gene-sets were downloaded from DisGeNET [75]. These include Alzheimer’s disease, epilepsy, and major psychiatric disorders (major depression, bipolar disorder, and schizophrenia) disease datasets. To avoid size related bias and improve the specificity of pathway profiles [76–78], we further restricted our analysis to diseases with gene-set size between 10 to 500 significant differentially expressed genes. Pairwise-overlap between disease-associated gene-sets was calculated using hypergeometric [79] gene-overlap package in R version 3.6.0.

### Gene Ontology (GO) analysis

Pathways affected in different astrocyte subtypes and diseases were determined using hypergeometric overlap analysis (HGA) with a background of 21,196 genes (default, gene-overlap package). The astrocyte-associated gene-sets and disease-associated gene-sets were tested against gene ontology (GO) pathways associated with Biological Process (GOBP), Molecular Function (GOMF), and Cellular Component (GOCC). Updated lists of GO-pathways were obtained from the Bader-lab (http://download.baderlab.org/EM_Genesets/). To compare enrichment of pathways across different astrocyte subtypes and diseases, the −log10(p-value) was used to generate the heatmap. To better identify the character of biological changes in the overlap results, a focused analysis of forty a-priori functional themes was performed. As described in our previous study [80], the pathways were filtered based on the parent-child association between GO-terms in our list of significant pathways (child-pathways) and hand-picked parent-pathways representing the a-priori theme from the GO-database.

### Density-Index

To quantitatively summarize how common (close to 1) or unique (close to 0) a theme is across different astrocyte subtypes and diseases; we deployed a previously developed density-index [81]. For a given ***r* x *c*** matrix of −log10(p-value), a density-score is obtained as:

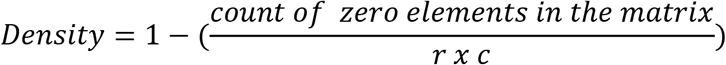

Where ***r*** and ***c*** represent the number of pathways in a theme and number of astrocyte subtypes and diseases, respectively. A zero element represents a non-significant disease-pathway relationship. For density associated with individual pathways and drug signatures, the density-index representing the fraction of non-zero elements in the number of astrocyte subtypes and diseases was used. In such an instance, the numerator of the above density formula represents count of zero elements in a vector. Whereas ***r*** and ***c*** in the denominator are constant holding a fixed value of 1 and 17, respectively.

### Drug-Target Enrichment Analysis

Enrichment of astrocyte subtypes and drug-induced molecular signatures in the disease-associated gene-sets was calculated using HGA. Drug-specific gene markers were downloaded from the DSigDB library of gene-sets [82]. In order to understand the druggable-mechanism and targets involved, gene-markers of drugs with known modes of action (MOA) and targets were used. Drugs were then further sorted by the typical diseases which the drug is used to treat.

### Rat Studies

Male Sprague-Dawley rats were housed with a 12-hour cycle of light and darkness. They were pair-housed and given access to food and water ad libitum. Rats were administered with either 28.5 mg kg^−1^ haloperidol-decanoate (n=10/group) via intramuscular injection every 3 weeks for 9 months or vehicle-treated with sesame oil (n=10/group) under the same conditions. At the end of the 9 months, brains were removed and stored at −80°C. All animal studies were carried out in accordance with the IACUC guidelines at the University of Alabama at Birmingham.

### qPCR Studies

RNA was extracted (RNeasy Mini Kit, QIAGEN) from 14μm thick fresh frozen sections of brain tissue as previously described [83, 84]. RNA was reverse-transcribed (High-Capacity cDNA Reverse Transcription Kit, Applied Biosystems) at cycling conditions: 1 cycle: 25°C for 10 minutes; 2 cycles: 37°C for 60 minutes; 1 cycle: 85°C for 5 minutes. cDNA was diluted and equalized to 11 ng/ul. Three microliters of cDNA and 17 μL of Taqman master mix (Applied Biosystems, Life Technologies) containing 1 μl of Taqman primer were loaded per well and ran at the following cycling conditions: 1 cycle: 95°C for 10 minutes; 40 cycles: 95°C for 15 seconds, 60°C for 60 seconds. Taqman primers for rats assayed were: GFAP (Rn01253033_m1), VIM (Rn00667825_m1), and SOX9 (Rn01751070_m1). The samples were run in duplicate. The relative gene expression was calculated by comparing to a standard curve made of a pool of cDNA from all samples. Gene expression was then normalized to the geomean of reference genes PPIA (Rn00690933_m1) and B2M (Rn00560865_m1). Data were analyzed for normal distribution. Data were log transformed and analyzed by Student’s t-test, p<0.05 for statistical tests. Data were analyzed with Graphpad Prism v7.04 (GraphPad Software, San Diego, CA, USA).

These commonly used astrocyte markers were selected because glial fibrillary acidic protein (GFAP) is highly expressed in many reactive astrocytes and is upregulated in response to neurological disorders [85]. Vimentin (VIM) is a marker in the cytoskeleton of astrocytes known to be upregulated in response to neurological injury and is also expressed by progenitor astrocytes [85]. Sex-determining region Y (SRY)-box 9 (SOX9) is a transcription factor whose expression also increases in response to disease or injury to astrocytes. [85].

### “Look up” studies

The R shiny application “Kaleidoscope” (https://kalganem.shinyapps.io/BrainDatabases/) [86] was used to search expression of GFAP, VIM, and SOX9 in publicly available transcriptomic datasets. The data is reported from analysis of rodent brain tissue following chronic (>2 weeks) administration with typical and atypical antipsychotics, antidepressants, and mood stabilizers. All individual dataset identifiers are listed in Table 5a. To determine the effects of antipsychotic medications on astrocyte marker gene expression in human brain, gene names GFAP, VIM, and SOX9 were searched in the Stanley Medical Research Institute (SMRI) Online Genomics Database repository [87, 88]. The SMRI contains postmortem brain transcriptomic data from patients diagnosed with schizophrenia and bipolar disorder who were either on or off antipsychotics and mood stabilizers, respectively, at time of death. The fold change and p-value of transcript expression in patients on/off medication is reported.

## Results

### Astrocyte subset genes most affected in neuropsychiatric disease

In order to identify a role for astrocyte subtypes in neuropsychiatric disease, we first assessed the enrichment of the gene-sets representing the five subtypes of astrocytes, described in methods and Batiuk et al. [74], and the gene-sets representing different psychiatric and neurological diseases from DisGeNET (Fig. 1).

**Figure 1.**
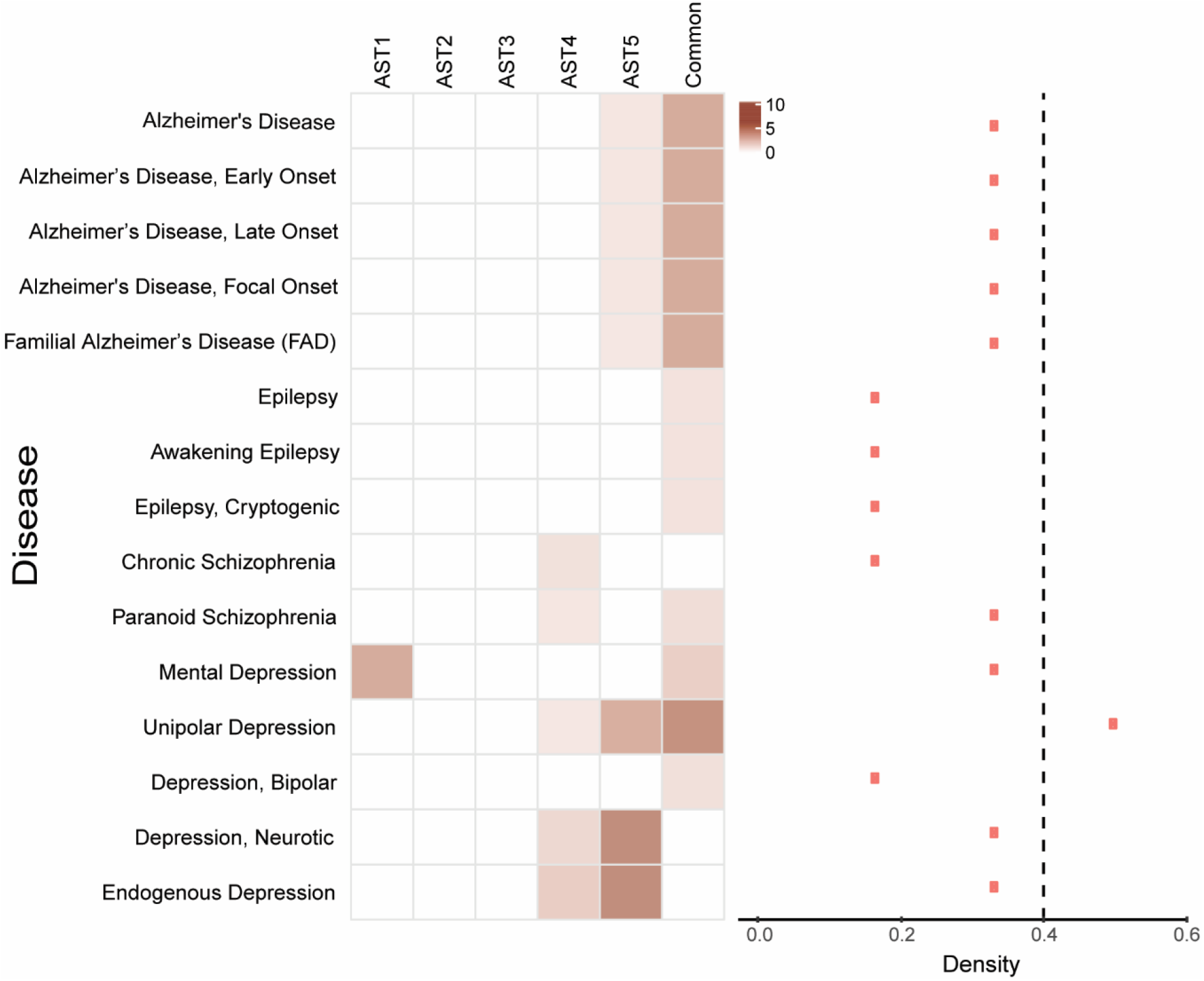
Heatmap displaying gene enrichment of unique subtypes of astrocytes across neuropsychiatric and neurological diseases. Color intensity is proportional to −log10 (p-value). Density index for all astrocyte subtypes across all disorders is displayed on right.

The set of genes representing the “common” astrocyte genes were the most frequently enriched across diseases, including Alzheimer’s disease, epilepsy, schizophrenia, and depression. This is in line with findings that astrocytes play important roles in the pathophysiology of these disorders [65]. The AST4 and AST5 subtypes, thought to represent progenitor astrocytes and intermediate (progenitor-mature) astrocyte populations, respectively [74], were also enriched across depression datasets. Additionally, AST4 genes were enriched in schizophrenia and AST5 genes were enriched in Alzheimer’s disease. Overall, unipolar depression is the disorder with the highest density index (>0.4). The density index quantitatively compares the significance of themes (disease gene expression) to an independent variable (astrocyte subtype) [81]. This suggests greater dysregulation of the astrocytes in this disorder compared to other neuropsychiatric disorders.

### Biological pathway enrichment across astrocyte subtypes and neuropsychiatric disease

In Figure 2, we explored the biological pathway gene enrichment of astrocyte subtypes and neurological and neuropsychiatric diseases using GOBP, GOMF and GOCC. Parent-child analysis was applied to cluster pathways [80]. First, looking at the enrichment between astrocyte subtypes and biological pathway clusters, we can see that gene enrichment of AST4 and AST5 subtypes is evident in “metabolic”, “plasticity/structure”, and “signaling” pathway clusters, which is in line with their expected functions as proposed in Batiuk et al. [74]. Gene enrichment of the “common” astrocyte subtype is found in similar clusters. Enrichment of the disease datasets shows “metabolic,” “signaling,” and “plasticity/structure” clusters are also enriched in Alzheimer’s disease and depression. Enrichment in the “plasticity/structure” cluster is also common to astrocyte subtypes and Alzheimer’s disease, depression, and epilepsy datasets. The pathway cluster enrichment seen in the schizophrenia datasets has limited overlap with enrichment seen in the astrocyte subtype groups, but the “immune” cluster is commonly enriched in the paranoid schizophrenia, Alzheimer’s disease, and depression datasets.

**Figure 2.**
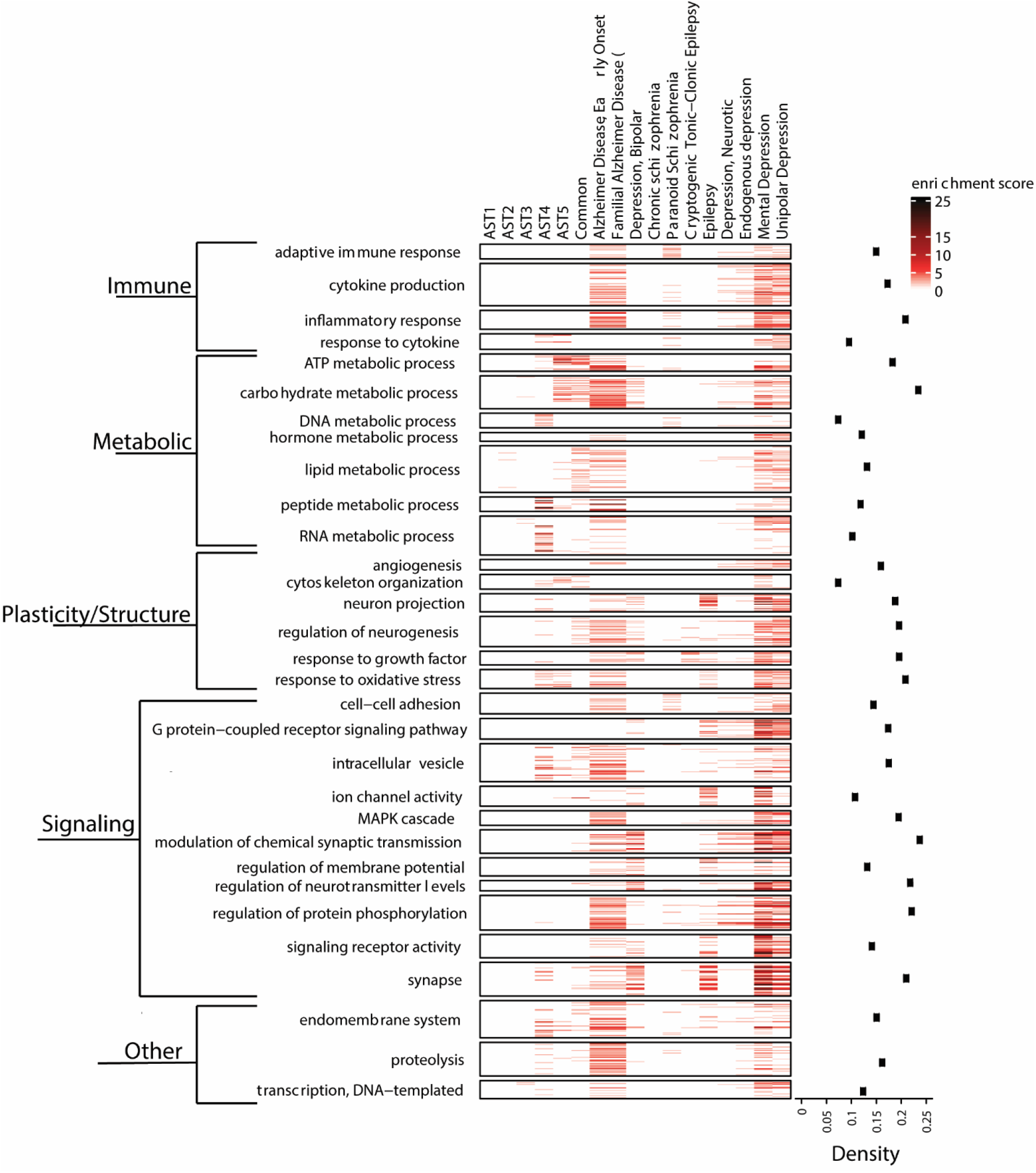
Heatmap displaying gene enrichment of clustered biological pathways across subtypes of astrocytes and neuropsychiatric and neurological diseases. Color intensity is proportional to −log10 (p-value). Density index for all pathways across astrocyte subtypes and disorders is displayed on right.

Specifically, common enriched pathways in astrocyte subtypes and across neuropsychiatric disease include ATP metabolism, carbohydrate metabolism, RNA metabolism, and peptide metabolism in the “metabolic” cluster, response to oxidative stress in the “plasticity/structure” pathway cluster, intracellular vesicle in the “signaling” cluster, and endomembrane system in the “other” cluster. These pathways broadly align with commonly dysregulated biological processes in CNS disorders, and may indicate a role for astrocytes in these pathological processes. The density index is highest in the plasticity/structure and signaling pathways, indicating that these pathways are the most commonly enriched across the astrocyte subtypes and neuropsychiatric disorders.

### Psychotropic medication effect on astrocyte subtypes and disease

Next, we assessed the gene enrichment of the astrocyte subtypes and neuropsychiatric diseases, and the gene-sets representing different types of commonly prescribed psychotropic medications for psychiatric and neurological disorders from the DSigDB library (Fig. 3). These include drug gene-sets for antipsychotics, antidepressants, mood stabilizers, Alzheimer’s medications, and other (adrenergic receptor antagonists and benzodiazepine receptor agonists) medications.

**Figure 3.**
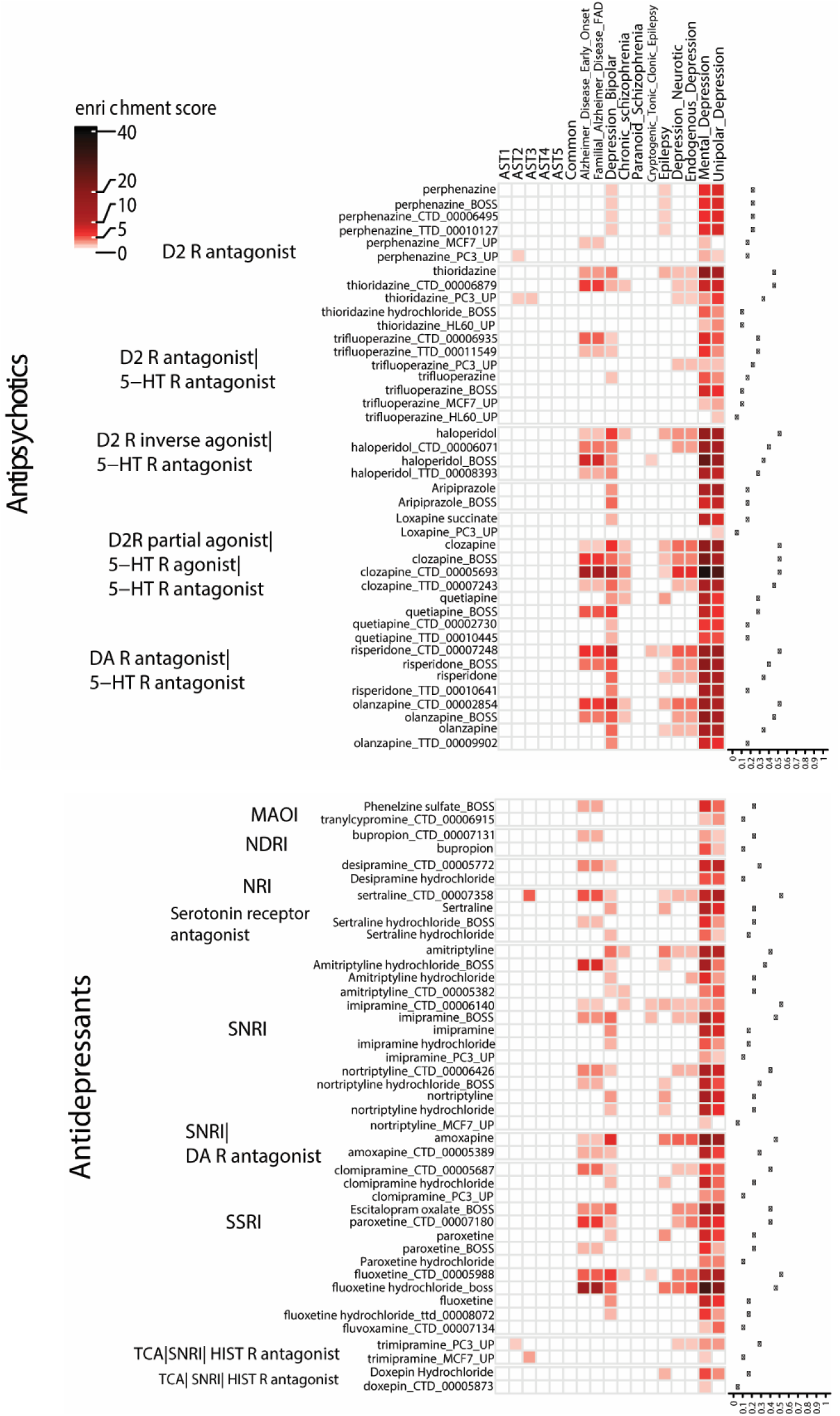
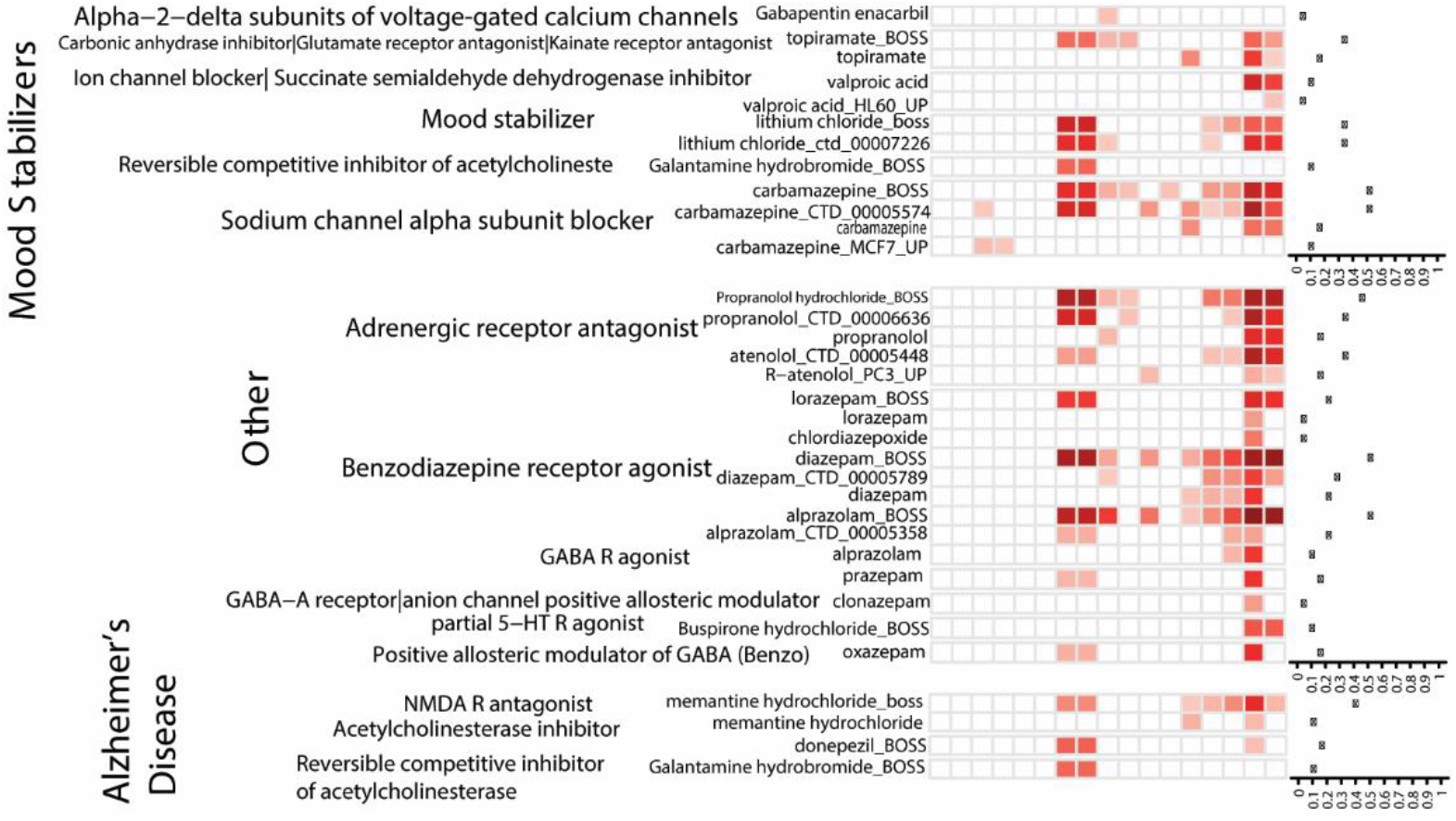
Heatmap displaying gene enrichment of different classes of psychotropic medications across astrocyte subtypes and neuropsychiatric and neurological diseases. Color intensity is proportional to −log10 (p-value). Density index for all pathways across astrocyte subtypes and disorders is displayed on right.

The first notable finding is that the overall gene enrichment of the astrocyte subtypes and psychotropic drug datasets is low. This is in contrast to the enrichment shown in the majority of the neuropsychiatric disorders and psychotropic drug datasets. Specifically, the Alzheimer’s disease and depression datasets showed high levels of enrichment with different medication groups. Other disease gene-sets, such as schizophrenia, showed low levels of enrichment across the medication types. It is worth mentioning that although many genes were enriched across disorders and the associated medication group (i.e. chronic schizophrenia and antipsychotics), genes were also enriched across non-linked disorders (i.e. unipolar depression and antipsychotics). Antipsychotics, including haloperidol, clozapine, risperidone, and olanzapine, and antidepressants, including imipramine and fluoxetine, had the highest density indices (>0.4) across analyses. This suggests that these drugs induce significant changes in gene expression that are common to gene expression associated with disease datasets but not astrocyte subtypes.

### Effect of chronic haloperidol medication on astrocyte marker expression in rat brain

In order to further explore the relationship between psychotropic medications and their effect on astrocytes, we assayed the mRNA levels of commonly used astrocyte markers GFAP, VIM, and SOX9 in the frontal cortex of rats treated chronically with the typical antipsychotic haloperidol-decanoate. There were no significant differences detected in the mRNA expression of GFAP (p=0.447), SOX9 (p=0.520), or VIM (p=0.778) in haloperidol-treated rats compared to vehicle-treated controls (p > 0.05; Fig. 4), suggesting that haloperidol does not significantly alter astrocyte gene marker expression.

**Figure 4.**
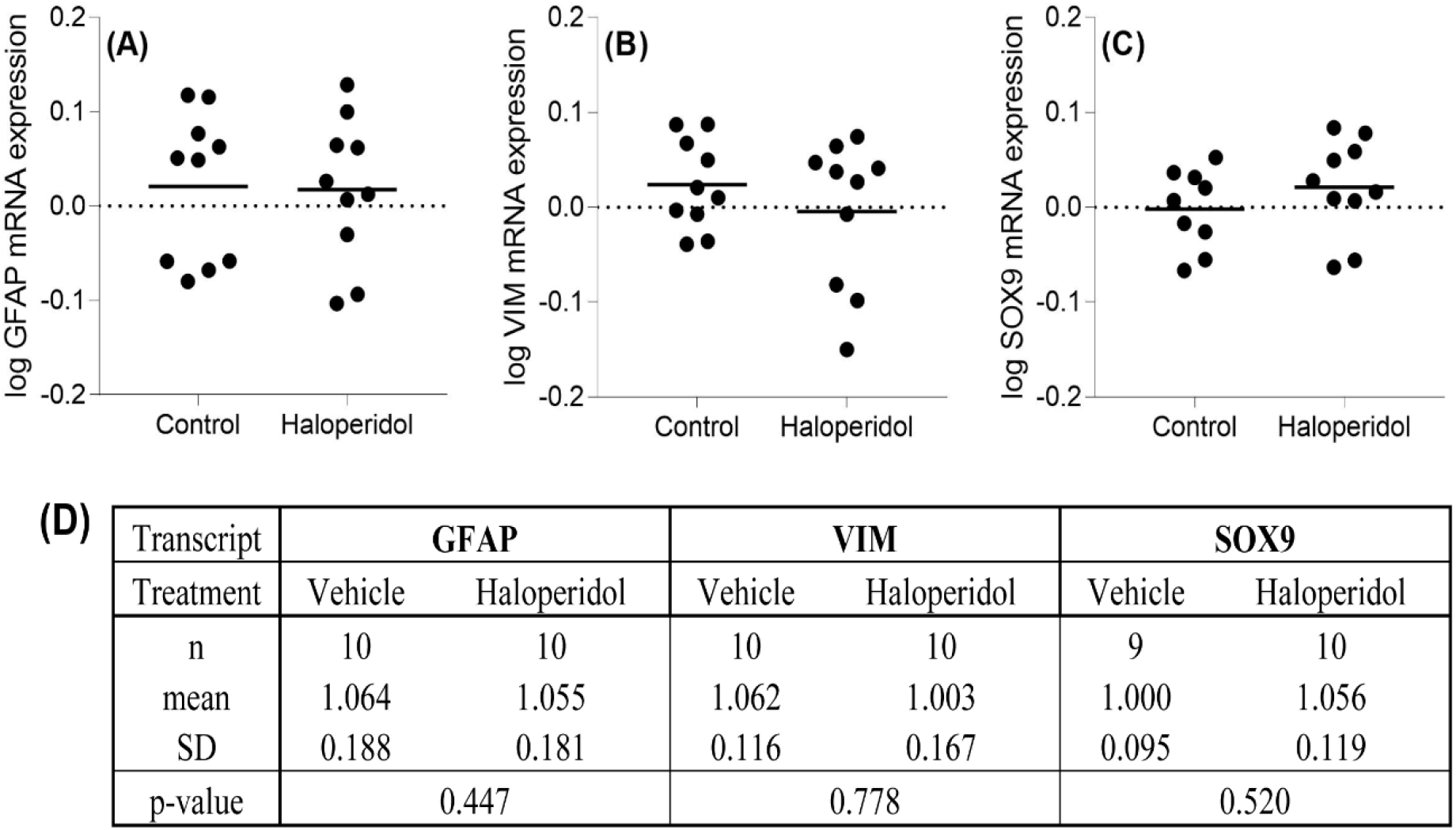
**(A-C)** Gene expression of astrocyte markers **(A)** GFAP, **(B)** VIM, and **(C)** SOX9 in frontal cortex of rats administered antipsychotic haloperidol-decanoate (n=10) or vehicle (n=lO) for 9 months. There was no significant difference (Student’s t-test, p>0.05) in transcript expression between any of the astrocyte markers in antipsychotic treated compared to vehicle. GFAP glial fibrillary acidic protein, VIM vimentin, SOX9 SRY (sex-determining region Y)-box 9. Data was log-transformed. Data expressed as mean. **(D)** Descriptive statistics of log-transformed gene expression of GFAP, VIM, and SOX9 in the haloperidol-treated rat studies. Data includes sample size of each group, the mean, standard deviation, and p-value comparing the vehicle-treated and haloperidol-treated groups. There were no significant differences in gene expression between vehicle and haloperidol groups in any of the markers using a significance level of 0.05.

### “Look up” Studies

To assess if other antipsychotic and psychotropic medications have a similar effect on astrocyte marker expression, we performed in silico “look-up” studies. In Figure 5a, we report log2 fold changes in astrocyte marker expression in rodents chronically treated with typical antipsychotics, atypical antipsychotics, antidepressants, and mood stabilizers. VIM and SOX9 were not significantly altered in any dataset. However, a subset of datasets (n=3) found significant changes in GFAP expression. These significant findings belong to treatment with an antidepressant, an atypical antipsychotic, and a typical antipsychotic, respectively, although the majority of data shows a lack of significant change as a result of psychotropic medication.

**Figure 5.**
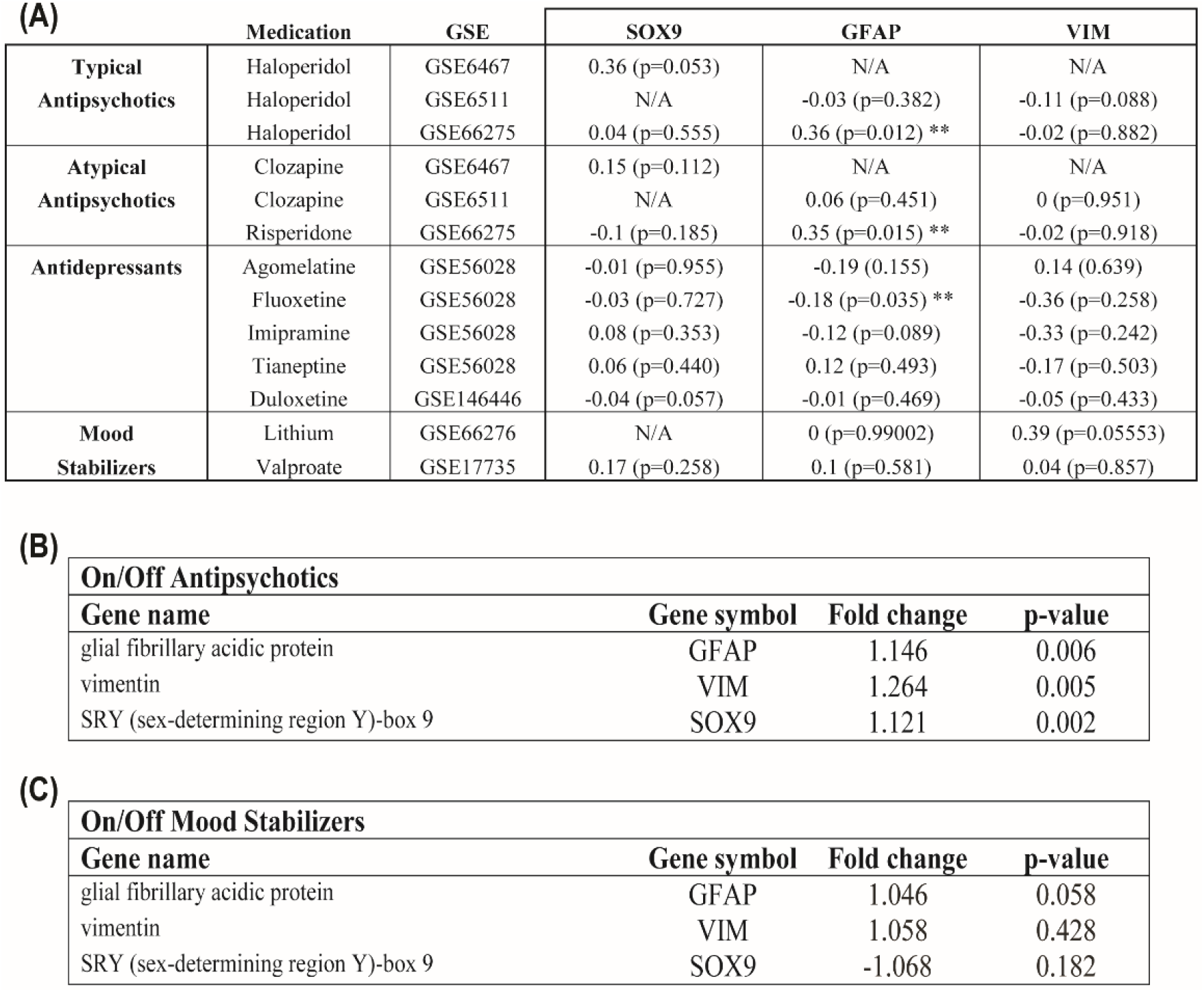
Look-up study report of differential gene expression of GFAP, VIM, and SOX9, in (A) rodent models of psychotropic drug administration and (B) postmortem brain tissue from schizophrenia and bipolar disorder subjects who are on/off psychotropic medication. A) L2FC expression and p-value for GFAP, VIM, and SOX9 expression in transcriptomic datasets from studies of rodent brain administered typical and atypical antipsychotics, antidepressants, or mood stabilizers for at least two weeks, were searched in Kaleidoscope. No significant changes in SOX9 or VIM expression were reported following administration of any psychotropic medications. A subset of datasets (n=3) reported significant (p<0.05) changes in GFAP expression after chronic administration of psychotropic medications. **B)** The fold change and p-value for transcript expression of GFAP, SOX9, and VIM in postmortem brain from schizophrenia subjects who were on versus off antipsychotics and bipolar disorder subjects who were on versus off mood stabilizers are reported. GFAP, VIM, and SOX9 transcript expression is significantly altered (p<0.05) in schizophrenia subjects who were on antipsychotics. There was no significant difference in transcript expression of any target in bipolar disorder subjects who were on mood stabilizers compared to those who were off medication. Data obtained from the Stanley Medical Research Institute (SMR1) Online Genomic database. GFAP glial fibrillary acidic protein, VIM vimentin, SOX9 SRY (sex-determining region Y)- box 9, FC fold change.

Next, we looked at the effects of medication on GFAP, VIM, and SOX9 mRNA expression in postmortem brain tissue from schizophrenia or bipolar subjects who are “on” and “off” medication, from the SMRI database (Fig. 5b-c). Expression of GFAP (FC=1.146, p=0.006), VIM (FC=1.264, p=0.005), and SOX9 (FC=1.121, p=0.002) were significantly increased (Fig. 5b) in schizophrenia subjects who were “on” antipsychotic medication at time of death. There was no significant difference in astrocyte marker transcript expression in GFAP (FC=1.046, p=0.058), VIM (FC=1.058, p=0.428), or SOX9 (FC=-1.068, p=0.182) (Fig. 5c), in bipolar subjects who were “on” mood stabilizers compared to subjects who were “off” medication at time of death.

## Discussion

Astrocytes play an important, albeit still poorly understood role in many neurological and psychiatric diseases [1–3, 65]. Although postmortem studies generally support altered astrocyte expression and morphology in neurological and psychiatric diseases [6, 7, 10, 20–29], many studies have shown conflicting findings [4–19] or report no changes in astrocytes in disease [27, 30–35].

The diversity of astrocyte subtypes [43, 45], their responses to different insults [43, 48, 61], and the potential effects of psychotropic medications on astrocytes [12, 15, 23-25, 67–73] may contribute to challenges in defining and interpreting astrocyte changes in the brain in these disorders.

To improve our understanding of how astrocytes are altered in the brain in disease and better our understanding of the effects of medications on astrocytes in the brain, we carried out bioinformatic analysis, taking advantage of emerging single-cell RNAseq datasets that define murine astrocyte subtypes, and human CNS disorder transcriptomic datasets. As unique molecular signatures for different subtypes of human astrocytes have yet to be elucidated [65, 66], we instead used signatures for five different astrocyte subtypes (AST1-5) and a “common” astrocyte signature derived from mouse brain [74].

First, we conducted gene enrichment analysis [81] to assess enrichment of different astrocyte subtypes (from Batiuk et al.) [74] in CNS disorders (from DSigDB) [82]. At least one subtype of astrocyte was enriched in every disease, with highest enrichment in “depression” disease datasets. The “common,” “AST4,” and “AST5” astrocyte gene signatures had the highest enrichment across disease types. AST4 and AST5 are developmental subtypes of astrocytes as opposed to the mature AST1-3 subtypes and may indicate a relationship between astrocyte cell immaturity and the formation of the reactive astrogliosis state [74].

Overall, enrichment of astrocyte subtypes in CNS disorders was relatively low except in depression, suggesting limited changes in gene expression of these cells in disease. However, species-specific differences in molecular signatures of astrocyte subtypes, and comparing astrocyte subtypes from “healthy” mice to human disease datasets [43, 48, 61] may contribute to low levels of enrichment. Although there is extensive conservation of gene expression between human and mouse astrocytes [89], there are also significant differences in the genes involved in responses to stressors such as inflammation [89], which could account for differences in astrocyte gene enrichment in human disorders.

Next, we looked at how astrocytes’ functions relate to disease and saw that many of the same biological pathways were enriched in murine astrocyte subtypes (“common” and “AST4-5”) and CNS disorders. Pathway analysis identified common enrichment in the “plasticity/structure” and “signaling” clusters, highlighting the role of astrocytes in these processes and the dysregulation of these processes in disease, especially Alzheimer’s disease and MDD [65, 90, 91]. Little enrichment of biological pathways was seen in the schizophrenia disease datasets, but this may reflect limitations of the available disease datasets rather than a lack of perturbation in biological pathways in this disorder.

Unsurprisingly, the pathway “response to oxidative stress” is enriched in astrocytes and across several diseases including Alzheimer’s disease, epilepsy, and depression. This is a proposed mechanism underlying the stress that causes astrocytes to become reactive in neuropsychiatric disorders [92], and therefore regulation of oxidative stress as a result of neurological disease is a key function of astrocytes. Human astrocytes have greater susceptibility to oxidative stress than mouse astrocytes [89], suggesting that enrichment may be even more pronounced in human astrocytes. Overall, this suggests a relationship between astrocyte functions and disease.

A potentially confounding variable in understanding the role of astrocytes in disease in postmortem studies is the effect of psychotropic medications that could be reversing disease-related changes in astrocytes at the structural, molecular, or functional level [23, 24, 26, 71, 73, 93, 94]. Postmortem studies that have considered the effects of psychotropic medications have found changes in astrocyte cell counts [71], morphology [73], and gene expression markers [15]. But it is not clear if these changes are a therapeutic result of medication use, an effect of the disease process, or a combination of both. Additionally, if astrocytes are altered in the disease state or as a result of medication, this may support their use as a therapeutic target for these diseases. To address these questions, we looked at enrichment of gene signatures induced by commonly prescribed psychotropic drugs in astrocyte subtypes and CNS disorder disease datasets. Overall, psychotropic drugs were highly enriched in disease datasets such as Alzheimer’s disease and depression, but there was surprisingly limited enrichment in astrocyte subtypes. This suggests that psychotropic drugs do not induce significant changes in gene expression of astrocyte subtypes.

While the general lack of drug enrichment in astrocyte subtypes is somewhat surprising [95, 96], it is possible that these medications do not affect astrocytes at the transcript level, or that there are inherent differences in the way that psychotropic medications affect astrocytes in different species and in the healthy versus diseased state as already discussed [43, 48, 61].

We also see enrichment of psychotropic drug gene sets in disorders for which those drugs are not commonly prescribed. For example, several classes of antipsychotics are enriched across depression datasets. This could be a consequence of the propensity of psychotropic medications to be “dirty” medications and have several off-target effects [97, 98]. This also may be related to the clinical practice of drug repurposing, or using medications in new or off-label ways to find alternative therapeutic benefits [99]. With drug repurposing, medications such as antipsychotics have been used off-label to treat other neuropsychiatric disorders even without the primary indication of the medication, such as psychotic symptoms. In clinical practice, this is most commonly used to treat various forms of depression, including major and vascular depression [100, 101]. Other bioinformatic approaches to drug repurposing have also shown enrichment of off-label psychotropic medications in non-associated neuropsychiatric conditions, identifying potential candidates for future drug repurposing in disorders such as schizophrenia, bipolar disorder, Alzheimer’s disease, and dementia [102, 103]. However, overall, these bioinformatic findings suggest that astrocytes are not significant targets of these psychotropic medications in neuropsychiatric disorders.

To confirm these findings from the gene enrichment analysis, we directly assessed the effects of chronic antipsychotic administration on astrocyte marker transcript expression in rodent brain. We also carried out additional “look-up” studies of rodent and postmortem transcriptomic brain analyses of the effects of antipsychotics, antidepressants, and mood stabilizers. In line with the bioinformatic analysis, transcript expression of astrocyte markers GFAP, vimentin, and SOX9 were not significantly altered in the rat frontal cortex following chronic administration of the antipsychotic haloperidol-decanoate.

Although there are limitations in using these markers, for example GFAP is not expressed in all astrocytes and is not always expressed only in astrocytes [104, 105], these are commonly used astrocyte markers with conserved expression across species, and they play differing, yet specific, roles in the reactive astrogliosis response in rats [106–109]. Taken altogether, these gene expression markers should provide a robust way to identify astrocytes and characterize astrocytic change, if present. Similar results were found in the “look-up” studies, which allow us to explore data sets that use different medication types and at different time-points. GFAP gene expression was significantly increased in a small subset of studies following antipsychotic administration. No significant changes were found in SOX9 or vimentin transcript expression following administration of any psychotropic drugs.

Overall, the “look-up” studies, qPCR analysis, and bioinformatics analysis suggest that antipsychotics have limited effect on transcript expression of glial markers in rodent models. Importantly, look-up studies of GFAP, vimentin, and SOX9 did find significant increases in expression of all gene targets in postmortem brain of schizophrenia subjects who were “on” antipsychotics compared to those who were “off” antipsychotics. No effect on marker expression was found in bipolar disorder subjects who were “on” mood stabilizers compared to those who were “off”. Data was not available for MDD subjects who were on/off antidepressant medications. This highlights the importance of considering specific disease- and drug-interaction effects on astrocytes as well as the limitations of studying rodent-derived astrocyte subtype molecular signatures.

## Limitations

A necessary limitation of this study is the use of astrocyte subtype molecular signatures derived from healthy mice for bioinformatic analysis, which may not reflect changes occurring in astrocytes in human disease or in response to psychotropic medication. Another limitation is the focus on transcript level expression changes. Many drugs act at the protein level rather than at the transcript level and their effects on astrocytes may not be captured by gene enrichment analysis.

## Conclusion and Future Directions

In summary, this study provides a unique analysis of archived transcriptomic data and emerging molecular characterization of astrocyte subtypes to illuminate the relationship between psychotropic medications and astrocytes in the context of neuropsychiatric disease. Surprisingly, our bioinformatics analysis suggests minimal enrichment of psychotropic medications in astrocyte subtypes, supported (generally) by “lookup” studies and qPCR in rodents. This suggests that these medications have limited effects on astrocytes or on inducing gene expression changes in astrocytes. However, transcript expression in postmortem studies for subjects on/off medications, suggests antipsychotics affect astrocyte expression, although this effect is not seen with mood stabilizers. This may contribute to conflicting findings in postmortem literature, if medications are reversing disease-related astrocyte expression changes. Additionally, given the relationships we found between important astrocyte functions and dysregulated processes in disease, it may be possible that reactive astrocytes contribute to the pathogenesis of disease and could serve as potential novel therapeutic targets in the future.

Our analysis is somewhat stymied by reliance on mouse astrocyte subtype molecular signatures from healthy brain tissue as equivalent human signatures are not available. This highlights the importance of conducting truly translational studies, including postmortem brain studies that account for the effects of medication. These findings serve as a starting point to better understanding the complex relationship between psychotropic medications and astrocytes in neuropsychiatric disorders, which could have significant consequences on our understanding of the pathophysiology of disease and potentially offer insight into targeting astrocytes therapeutically.

## References

1. Pekny, M., et al., Astrocytes: a central element in neurological diseases. Acta Neuropathol, 2016. 131(3): p. 323–45.

2. Kettenmann, H., B.R. Ransom, and Oxford University Press., Neuroglia. 2013, Oxford University Press,: New York. p. 1 online resource (xxii, 930 p.

3. Chen, Y., et al., The role of astrocytes in oxidative stress of central nervous system: A mixed blessing. Cell Prolif, 2020. 53(3): p. e12781.

4. Toro, C.T., et al., Glial fibrillary acidic protein and glutamine synthetase in subregions of prefrontal cortex in schizophrenia and mood disorder. Neurosci Lett, 2006. 404(3): p. 276–81.

5. Markova, E., et al., 3-D Golgi and image analysis of the olfactory tubercle in schizophrenia. Anal Quant Cytol Histol, 2000. 22(2): p. 178–82.

6. Feresten, A.H., et al., Increased expression of glial fibrillary acidic protein in prefrontal cortex in psychotic illness. Schizophr Res, 2013. 150(1): p. 252–7.

7. Qi, X.R., W. Kamphuis, and L. Shan, Astrocyte Changes in the Prefrontal Cortex From Aged Non-suicidal Depressed Patients. Front Cell Neurosci, 2019. 13: p. 503.

8. Rao, J.S., et al., Increased excitotoxicity and neuroinflammatory markers in postmortem frontal cortex from bipolar disorder patients. Mol Psychiatry, 2010. 15(4): p. 384–92.

9. Williams, M.R., et al., Astrocyte decrease in the subgenual cingulate and callosal genu in schizophrenia. Eur Arch Psychiatry Clin Neurosci, 2013. 263(1): p. 41–52.

10. Williams, M., et al., Fibrillary astrocytes are decreased in the subgenual cingulate in schizophrenia. Eur Arch Psychiatry Clin Neurosci, 2014. 264(4): p. 357–62.

11. Altshuler, L.L., et al., Amygdala astrocyte reduction in subjects with major depressive disorder but not bipolar disorder. Bipolar Disord, 2010. 12(5): p. 541–9.

12. Torres-Platas, S.G., et al., Glial fibrillary acidic protein is differentially expressed across cortical and subcortical regions in healthy brains and downregulated in the thalamus and caudate nucleus of depressed suicides. Mol Psychiatry, 2016. 21(4): p. 509–15.

13. Choudary, P.V., et al., Altered cortical glutamatergic and GABAergic signal transmission with glial involvement in depression. Proc Natl Acad Sci U S A, 2005. 102(43): p. 15653–8.

14. Zhao, J., et al., Prefrontal changes in the glutamate-glutamine cycle and neuronal/glial glutamate transporters in depression with and without suicide. J Psychiatr Res, 2016. 82: p. 8–15.

15. Fatemi, S.H., et al., Glial fibrillary acidic protein is reduced in cerebellum of subjects with major depression, but not schizophrenia. Schizophr Res, 2004. 69(2-3): p. 317–23.

16. Arnold, S.E., et al., Glial fibrillary acidic protein-immunoreactive astrocytosis in elderly patients with schizophrenia and dementia. Acta Neuropathol, 1996. 91(3): p. 269–77.

17. Muller, M.B., et al., Neither major depression nor glucocorticoid treatment affects the cellular integrity of the human hippocampus. Eur J Neurosci, 2001. 14(10): p. 1603–12.

18. Hercher, C., V. Chopra, and C.L. Beasley, Evidence for morphological alterations in prefrontal white matter glia in schizophrenia and bipolar disorder. J Psychiatry Neurosci, 2014. 39(6): p. 376–85.

19. Johnston-Wilson, N.L., et al., Disease-specific alterations in frontal cortex brain proteins in schizophrenia, bipolar disorder, and major depressive disorder. The Stanley Neuropathology Consortium. Mol Psychiatry, 2000. 5(2): p. 142–9.

20. Pakkenberg, B., Pronounced reduction of total neuron number in mediodorsal thalamic nucleus and nucleus accumbens in schizophrenics. Arch Gen Psychiatry, 1990. 47(11): p. 1023–8.

21. Barley, K., S. Dracheva, and W. Byne, Subcortical oligodendrocyte- and astrocyte-associated gene expression in subjects with schizophrenia, major depression and bipolar disorder. Schizophr Res, 2009. 112(1-3): p. 54–64.

22. Bowley, M.P., et al., Low glial numbers in the amygdala in major depressive disorder. Biol Psychiatry, 2002. 52(5): p. 404–12.

23. Ongur, D., W.C. Drevets, and J.L. Price, Glial reduction in the subgenual prefrontal cortex in mood disorders. Proc Natl Acad Sci U S A, 1998. 95(22): p. 13290–5.

24. Rajkowska, G., et al., Morphometric evidence for neuronal and glial prefrontal cell pathology in major depression. Biol Psychiatry, 1999. 45(9): p. 1085–98.

25. Cotter, D., et al., Reduced neuronal size and glial cell density in area 9 of the dorsolateral prefrontal cortex in subjects with major depressive disorder. Cereb Cortex, 2002. 12(4): p. 386–94.

26. Cotter, D., et al., Reduced glial cell density and neuronal size in the anterior cingulate cortex in major depressive disorder. Arch Gen Psychiatry, 2001. 58(6): p. 545–53.

27. Catts, V.S., et al., Increased expression of astrocyte markers in schizophrenia: Association with neuroinflammation. Aust N Z J Psychiatry, 2014. 48(8): p. 722–34.

28. Torres-Platas, S.G., et al., Astrocytic hypertrophy in anterior cingulate white matter of depressed suicides. Neuropsychopharmacology, 2011. 36(13): p. 2650–8.

29. Strakowski, S.M., et al., The functional neuroanatomy of bipolar disorder: a consensus model. Bipolar Disord, 2012. 14(4): p. 313–25.

30. Steffek, A.E., et al., Cortical expression of glial fibrillary acidic protein and glutamine synthetase is decreased in schizophrenia. Schizophr Res, 2008. 103(1-3): p. 71–82.

31. Trepanier, M.O., et al., Postmortem evidence of cerebral inflammation in schizophrenia: a systematic review. Mol Psychiatry, 2016. 21(8): p. 1009–26.

32. Pantazopoulos, H., et al., Extracellular matrix-glial abnormalities in the amygdala and entorhinal cortex of subjects diagnosed with schizophrenia. Arch Gen Psychiatry, 2010. 67(2): p. 155–66.

33. Damadzic, R., et al., A quantitative immunohistochemical study of astrocytes in the entorhinal cortex in schizophrenia, bipolar disorder and major depression: absence of significant astrocytosis. Brain Res Bull, 2001. 55(5): p. 611–8.

34. Dean, B., L. Gray, and E. Scarr, Regionally specific changes in levels of cortical S100beta in bipolar 1 disorder but not schizophrenia. Aust N Z J Psychiatry, 2006. 40(3): p. 217–24.

35. Webster, M.J., et al., Immunohistochemical localization of phosphorylated glial fibrillary acidic protein in the prefrontal cortex and hippocampus from patients with schizophrenia, bipolar disorder, and depression. Brain Behav Immun, 2001. 15(4): p. 388–400.

36. Benes, F.M., J. Davidson, and E.D. Bird, Quantitative cytoarchitectural studies of the cerebral cortex of schizophrenics. Arch Gen Psychiatry, 1986. 43(1): p. 31–5.

37. Bruton, C.J., et al., Schizophrenia and the brain: a prospective clinico-neuropathological study. Psychol Med, 1990. 20(2): p. 285–304.

38. Rajkowska, G., et al., Layer-specific reductions in GFAP-reactive astroglia in the dorsolateral prefrontal cortex in schizophrenia. Schizophr Res, 2002. 57(2-3): p. 127–38.

39. Schmitt, A., et al., Stereologic investigation of the posterior part of the hippocampus in schizophrenia. Acta Neuropathol, 2009. 117(4): p. 395–407.

40. Verkhratsky, A., R. Zorec, and V. Parpura, Stratification of astrocytes in healthy and diseased brain. Brain Pathol, 2017. 27(5): p. 629–644.

41. Verkhratsky, A., J.J. Rodriguez, and L. Steardo, Astrogliopathology: a central element of neuropsychiatric diseases? Neuroscientist, 2014. 20(6): p. 576–88.

42. Verkhratsky, A., et al., Astroglial asthenia and loss of function, rather than reactivity, contribute to the ageing of the brain. Pflugers Arch, 2021. 473(5): p. 753–774.

43. Zamanian, J.L., et al., Genomic analysis of reactive astrogliosis. J Neurosci, 2012. 32(18): p. 6391–410.

44. Anderson, M.A., et al., Astrocyte scar formation aids central nervous system axon regeneration. Nature, 2016. 532(7598): p. 195–200.

45. Sofroniew, M.V., Molecular dissection of reactive astrogliosis and glial scar formation. Trends Neurosci, 2009. 32(12): p. 638–47.

46. Eddleston, M. and L. Mucke, Molecular profile of reactive astrocytes--implications for their role in neurologic disease. Neuroscience, 1993. 54(1): p. 15–36.

47. Pekny, M. and M. Nilsson, Astrocyte activation and reactive gliosis. Glia, 2005. 50(4): p. 427–34.

48. Wilhelmsson, U., et al., Redefining the concept of reactive astrocytes as cells that remain within their unique domains upon reaction to injury. Proc Natl Acad Sci U S A, 2006. 103(46): p. 17513–8.

49. Sofroniew, M.V., Reactive astrocytes in neural repair and protection. Neuroscientist, 2005. 11(5): p. 400–7.

50. Maragakis, N.J. and J.D. Rothstein, Mechanisms of Disease: astrocytes in neurodegenerative disease. Nat Clin Pract Neurol, 2006. 2(12): p. 679–89.

51. Correa-Cerro, L.S. and J.W. Mandell, Molecular mechanisms of astrogliosis: new approaches with mouse genetics. J Neuropathol Exp Neurol, 2007. 66(3): p. 169–76.

52. Zador, Z., et al., Role of aquaporin-4 in cerebral edema and stroke. Handb Exp Pharmacol, 2009(190): p. 159–70.

53. Simard, M. and M. Nedergaard, The neurobiology of glia in the context of water and ion homeostasis. Neuroscience, 2004. 129(4): p. 877–96.

54. Swanson, R.A., W. Ying, and T.M. Kauppinen, Astrocyte influences on ischemic neuronal death. Curr Mol Med, 2004. 4(2): p. 193–205.

55. Chen, Y., et al., Astrocytes protect neurons from nitric oxide toxicity by a glutathione-dependent mechanism. J Neurochem, 2001. 77(6): p. 1601–10.

56. Hamby, M.E., J.A. Hewett, and S.J. Hewett, TGF-beta1 potentiates astrocytic nitric oxide production by expanding the population of astrocytes that express NOS-2. Glia, 2006. 54(6): p. 566–77.

57. Christopherson, K.S., et al., Thrombospondins are astrocyte-secreted proteins that promote CNS synaptogenesis. Cell, 2005. 120(3): p. 421–33.

58. Stevens, B., et al., The classical complement cascade mediates CNS synapse elimination. Cell, 2007. 131(6): p. 1164–78.

59. Burda, J.E. and M.V. Sofroniew, Reactive gliosis and the multicellular response to CNS damage and disease. Neuron, 2014. 81(2): p. 229–48.

60. Sofroniew, M.V. and H.V. Vinters, Astrocytes: biology and pathology. Acta Neuropathol, 2010. 119(1): p. 7–35.

61. Sofroniew, M.V., Astrogliosis. Cold Spring Harb Perspect Biol, 2014. 7(2): p. a020420.

62. Verkhratsky, A., et al., Astroglial atrophy in Alzheimer’s disease. Pflugers Arch, 2019. 471(10): p. 1247–1261.

63. Middeldorp, J. and E.M. Hol, GFAP in health and disease. Prog Neurobiol, 2011. 93(3): p. 421–43.

64. Norton, W.T., et al., Quantitative aspects of reactive gliosis: a review. Neurochem Res, 1992. 17(9): p. 877–85.

65. Zhang, X., et al., Role of Astrocytes in Major Neuropsychiatric Disorders. Neurochem Res, 2021.

66. Zhang, Y., et al., Purification and Characterization of Progenitor and Mature Human Astrocytes Reveals Transcriptional and Functional Differences with Mouse. Neuron, 2016. 89(1): p. 37–53.

67. Ho, B.C., et al., Long-term antipsychotic treatment and brain volumes: a longitudinal study of first-episode schizophrenia. Arch Gen Psychiatry, 2011. 68(2): p. 128–37.

68. Lieberman, J.A., et al., Antipsychotic drug effects on brain morphology in first-episode psychosis. Arch Gen Psychiatry, 2005. 62(4): p. 361–70.

69. Konopaske, G.T., et al., Effect of chronic antipsychotic exposure on astrocyte and oligodendrocyte numbers in macaque monkeys. Biol Psychiatry, 2008. 63(8): p. 759–65.

70. Konopaske, G.T., et al., Effect of chronic exposure to antipsychotic medication on cell numbers in the parietal cortex of macaque monkeys. Neuropsychopharmacology, 2007. 32(6): p. 1216–23.

71. Martin, J.L., P.J. Magistretti, and I. Allaman, Regulation of neurotrophic factors and energy metabolism by antidepressants in astrocytes. Curr Drug Targets, 2013. 14(11): p. 1308–21.

72. Miguel-Hidalgo, J.J., et al., Glial and glutamatergic markers in depression, alcoholism, and their comorbidity. J Affect Disord, 2010. 127(1-3): p. 230–40.

73. Rajkowska, G., et al., Differential effect of lithium on cell number in the hippocampus and prefrontal cortex in adult mice: a stereological study. Bipolar Disord, 2016. 18(1): p. 41–51.

74. Batiuk, M.Y., et al., Identification of region-specific astrocyte subtypes at single cell resolution. Nat Commun, 2020. 11(1): p. 1220.

75. Pinero, J., et al., The DisGeNET knowledge platform for disease genomics: 2019 update. Nucleic Acids Res, 2020. 48(D1): p. D845–D855.

76. Williams, A.G., et al., RNA-seq Data: Challenges in and Recommendations for Experimental Design and Analysis. Curr Protoc Hum Genet, 2014. 83: p. 11 13 1–20.

77. Holmans, P., Statistical methods for pathway analysis of genome-wide data for association with complex genetic traits. Adv Genet, 2010. 72: p. 141–79.

78. Wang, K., M. Li, and H. Hakonarson, Analysing biological pathways in genome-wide association studies. Nat Rev Genet, 2010. 11(12): p. 843–54.

79. Fuxman Bass, J.I., et al., Using networks to measure similarity between genes: association index selection. Nat Methods, 2013. 10(12): p. 1169–76.

80. Shukla, R., et al., The Relative Contributions of Cell-Dependent Cortical Microcircuit Aging to Cognition and Anxiety. Biol Psychiatry, 2019. 85(3): p. 257–267.

81. Smail, M.A., et al., Similarities and dissimilarities between psychiatric cluster disorders. Mol Psychiatry, 2021.

82. Yoo, M., et al., DSigDB: drug signatures database for gene set analysis. Bioinformatics, 2015. 31(18): p. 3069–71.

83. O’Donovan, S.M., et al., Glutamate transporter splice variant expression in an enriched pyramidal cell population in schizophrenia. Transl Psychiatry, 2015. 5: p. e579.

84. O’Donovan, S.M., et al., Cell-subtype-specific changes in adenosine pathways in schizophrenia. Neuropsychopharmacology, 2018. 43(8): p. 1667–1674.

85. Escartin, C., et al., Reactive astrocyte nomenclature, definitions, and future directions. Nat Neurosci, 2021. 24(3): p. 312–325.

86. Alganem, K., et al., Kaleidoscope: A New Bioinformatics Pipeline Web Application for In Silico Hypothesis Exploration of Omics Signatures. bioRxiv, 2020: p. 2020.05.01.070805.

87. Torrey, E.F., et al., The stanley foundation brain collection and neuropathology consortium. Schizophr Res, 2000. 44(2): p. 151–5.

88. Higgs, B.W., et al., An online database for brain disease research. BMC Genomics, 2006. 7: p. 70.

89. Li, J., et al., Conservation and divergence of vulnerability and responses to stressors between human and mouse astrocytes. Nat Commun, 2021. 12(1): p. 3958.

90. Verkhratsky, A., et al., Astroglia in Alzheimer’s Disease. Adv Exp Med Biol, 2019. 1175: p. 273–324.

91. Dossi, E., F. Vasile, and N. Rouach, Human astrocytes in the diseased brain. Brain Res Bull, 2018. 136: p. 139–156.

92. Rizor, A., et al., Astrocytic Oxidative/Nitrosative Stress Contributes to Parkinson’s Disease Pathogenesis: The Dual Role of Reactive Astrocytes. Antioxidants (Basel), 2019. 8(8).

93. Peng, L., B. Li, and A. Verkhratsky, Targeting astrocytes in bipolar disorder. Expert Rev Neurother, 2016. 16(6): p. 649–57.

94. Rivera, A.D. and A.M. Butt, Astrocytes are direct cellular targets of lithium treatment: novel roles for lysyl oxidase and peroxisome-proliferator activated receptor-gamma as astroglial targets of lithium. Transl Psychiatry, 2019. 9(1): p. 211.

95. Sethi, P., et al., Automated morphometric analysis with SMorph software reveals plasticity induced by antidepressant therapy in hippocampal astrocytes. J Cell Sci, 2021. 134(12).

96. Bouvier, M.L., et al., Sex-dependent alterations of dopamine receptor and glucose transporter density in rat hypothalamus under long-term clozapine and haloperidol medication. Brain Behav, 2020. 10(8): p. e01694.

97. Schulz, P. and T. Steimer, Psychotropic medication, psychiatric disorders, and higher brain functions. Dialogues Clin Neurosci, 2000. 2(3): p. 177–82.

98. Ericson, E., et al., Off-target effects of psychoactive drugs revealed by genome-wide assays in yeast. PLoS Genet, 2008. 4(8): p. e1000151.

99. Pushpakom, S., et al., Drug repurposing: progress, challenges and recommendations. Nat Rev Drug Discov, 2019. 18(1): p. 41–58.

100. Mohammad Sadeghi, H., et al., Drug Repurposing for the Management of Depression: Where Do We Stand Currently? Life (Basel), 2021. 11(8).

101. Ebada, M.E., Drug repurposing may generate novel approaches to treating depression. J Pharm Pharmacol, 2017. 69(11): p. 1428–1436.

102. So, H.C., et al., Translating GWAS findings into therapies for depression and anxiety disorders: gene-set analyses reveal enrichment of psychiatric drug classes and implications for drug repositioning. Psychol Med, 2019. 49(16): p. 2692–2708.

103. Luscher Dias, T., et al., Drug repositioning for psychiatric and neurological disorders through a network medicine approach. Transl Psychiatry, 2020. 10(1): p. 141.

104. Testen, A., R. Kim, and K.J. Reissner, High-Resolution Three-Dimensional Imaging of Individual Astrocytes Using Confocal Microscopy. Curr Protoc Neurosci, 2020. 91(1): p. e92.

105. Kimelberg, H.K., The problem of astrocyte identity. Neurochem Int, 2004. 45(2-3): p. 191–202.

106. O’Leary, L.A., et al., Characterization of Vimentin-Immunoreactive Astrocytes in the Human Brain. Front Neuroanat, 2020. 14: p. 31.

107. Quist, E., H. Ahlenius, and I. Canals, Transcription Factor Programming of Human Pluripotent Stem Cells to Functionally Mature Astrocytes for Monocultures and Cocultures with Neurons. Methods Mol Biol, 2021. 2352: p. 133–148.

108. Balouch, B., et al., Conventional immunomarkers stain a fraction of astrocytes in vitro: A comparison of rat cortical and spinal cord astrocytes in naive and stimulated cultures. J Neurosci Res, 2021. 99(3): p. 806–826.

109. Dragic, M., et al., Two Distinct Hippocampal Astrocyte Morphotypes Reveal Subfield-Different Fate during Neurodegeneration Induced by Trimethyltin Intoxication. Neuroscience, 2019. 423: p. 3854.

